# Molecular determinants of enterotoxigenic *Escherichia coli* heat-stable toxin secretion and delivery

**DOI:** 10.1101/299313

**Authors:** Yuehui Zhu, Qingwei Luo, Sierra M. Davis, Chase Westra, Tim J. Vickers, James M. Fleckenstein

## Abstract

Enterotoxigenic *Escherichia coli* (ETEC), a heterogeneous diarrheal pathovar defined by production of heat-labile (LT) and/or heat-stable (ST) toxins, remain major causes of mortality among children in developing regions, and cause substantial morbidity in individuals living in or traveling to endemic areas. Studies demonstrating a major burden of ST-producing ETEC have focused interest on ST toxoids for ETEC vaccines. We therefore examined fundamental aspects of ETEC ST biology using ETEC H10407, which carries *estH* and *estP* genes encoding ST-H and ST-P, respectively, in addition to *eltAB* genes responsible for LT. In this background, we found that deletion of *estH* significantly diminished cGMP activation in target epithelia, while deletion of *estP* had a surprisingly modest impact, and a dual *estH/estP* mutant was not appreciably different than the *estH* mutant. Nevertheless, either ST-H or ST-P recombinant peptides stimulated cGMP production. We also found that the TolC efflux protein was essential for both toxin secretion and delivery, providing a potential avenue for efflux inhibitors in treatment of acute diarrheal illness. In addition, we demonstrated that the EtpA adhesin is required for optimal delivery of ST and that antibodies against either the adhesin or ST-H significantly impaired toxin delivery and cGMP activation in target T84 cells. Finally, we used FLAG epitope fusions to demonstrate that the ST-H pro-peptide sequence is secreted by the bacteria, potentially providing additional targets for antibody neutralization. These studies collectively extend our understanding of ETEC pathogenesis and potentially inform additional avenues to mitigate disease by these common diarrheal pathogens.

## Introduction

Diarrheal illnesses in low income countries continue to cause substantial morbidity and remain one of the leading causes of death in young children in developing countries under the age of five years. Among the bacterial causes of diarrheal illness enteroxigenic *Escherichia coli* (ETEC) are commonly linked to more severe forms of illness in young children(<u>1</u>). These organisms are perennially the most common cause of diarrhea in those who travel to endemic areas where sanitation is poor(<u>2</u>, <u>3</u>), however they have been identified repeatedly as the etiology of diarrheal outbreaks and sporadic cases of illness in industrialized countries including the U.S(<u>4–8</u>).

Acute clinical presentations of ETEC infection may range from mild self-limited illness to severe cholera-like diarrhea(<u>9–11</u>). In addition, ETEC and other diarrheal pathogens have been linked to pernicious sequelae of malnutrition, growth stunting, and impaired cognitive(<u>12</u>) development. Presently, there are no vaccines to protect against these common infections. ETEC are a genetically(<u>13</u>) and serotypically (<u>14</u>) diverse pathovar of *E. coli* defined by the production of heat-labile (LT) and/or heat stable (ST) enterotoxins which activate production of host cyclic nucleotides to alter intestinal salt and water transport that culminate in net fluid losses and diarrhea.

Heat-stable toxins are synthesized as 72 amino acid proteins consisting of a signal peptide, a pro peptide and a carboxy terminal region of 18-19 amino acids, which forms the mature active enterotoxin(<u>15</u>). Two enterotoxins cause disease in humans: STp (ST1a) 18 amino acids, and STh (ST1b) 19 amino acids. Both mature toxins contain four cysteine residues which form two intramolecular disulfide bonds. Their overall structure is shared with two homologous mammalian peptides, guanylin and uroguanylin. Each of the bacterial and mammalian peptides binds to guanylate cyclase C(<u>16</u>, <u>17</u>), leading to increases in intracellular cGMP(<u>18</u>). Increases in this cyclic nucleotide result in activation of protein kinases which phosphorylate and activate the cystic fibrosis transmembrane regulatory(CFTR) channel, and inhibit sodium-hydrogen ion exchange(<u>19</u>). These effects lead to a net loss of salt and water into the intestinal lumen with ensuing watery diarrhea.

Bacteria producing any of the toxins LT, STh, or STp have been linked to diarrheal illness in humans(<u>20–23</u>), and recent studies suggest that ST-producing ETEC are commonly represented among the pathogens that cause severe diarrheal illness among young children in low income countries leading to substantial interest in the development of a vaccine that incorporates ST-toxoids(<u>24</u>).

Enterotoxigenic *E. coli* strain H10407, originally isolated from a case of severe cholera-like diarrheal illness in Bangladesh, is to date the most extensively characterized isolate of this pathovar. Interestingly, this isolate encodes all three enterotoxins associated with ETEC diarrheal illness in humans(<u>25</u>, <u>26</u>), with the gene for STh on the largest 94,797 bp virulence plasmid (<u>NCBI Genbank acccession NC_017724.1)</u>, and the genes for both LT and STp on a 66681 bp plasmid(<u>27</u>). H10407 is frequently used as the challenge strain in controlled human infection models to test candidate vaccines. Therefore, we set out to examine the relative contribution of STh and STp to activation of cGMP in host epithelia by H10407, and the ability of anti-ST and anti-adhesin antibodies to mitigate effective toxin delivery by the bacteria.

## Materials and methods

### Bacterial strains and growth conditions

Bacterial strains used are listed in Table 1. For general purposes, ETEC bacteria were grown in lysogeny broth at 37°C overnight from frozen glycerol stocks. To optimize synthesis and secretion of ST toxin, bacteria were grown in Casamino Acids-yeast extract –sucrose medium (CAYE-ST) (2% Casamino acids, 0.6% Yeast extract, 43mM NaCl, 38mM K_2_HPO_4_, 203mM MgSO_4_, 25mM MnCl_2_, 18mM FeCl_3_, 2% sucrose) at 37°C overnight from frozen glycerol stocks.

**Table 1.**

| table 1 bacterial strains and plasmids used in these studies |  |  |
| --- | --- | --- |
| Strain or plasmid | Description | Source or reference(s) |
| Strains |  |  |
| H10407 | Wild-type ETEC strain O78:H11; CFA/1; LT, ST-H, ST-P; EtpA | (27, 37, 53) |
| jf2656 | H10407 derivative with isogenic deletion of *estP* | This study |
| jf2649 | H10407 derivative with isogenic deletion of *estH* | This study |
| jf2847 | H10407 derivative with isogenic deletion of *estP* and *estH* | This study |
| jf3038 | deletion of *astA* in on the large p948 virulence plasmid of H10407; Km<sup>R</sup> | this study |
| jf3081 | jf2649 complemented with pQL231 | this study |
| JW5503-1 | *E. coli* K-12 in-frame ΔtolC732:: Km<sup>R</sup> | (30) |
| jf4652 | H10407 derivative with tolC::Km<sup>R</sup> mutation | this study |
| jf4709 | jf4652 with pBAD/mycHisB | this study |
| jf4712 | jf4652 complemented with pYZ008 | this study |
| jf4644 | jf2847 with pFLAG-CTC, Cm<sup>R</sup> Km<sup>R</sup> Amp<sup>R</sup> | this study |
| jf4648 | jf2847 with pFLAG-STh, Cm<sup>R</sup> Km<sup>R</sup> Amp<sup>R</sup> | this study |
| jf4651 | jf2847 with pSTh-FLAG, Cm<sup>R</sup> Km<sup>R</sup> Amp<sup>R</sup> | this study |
| AAEC191A | afimbriate *E. coli* K-12 derivative | (33) |
| Plasmids |  |  |
| pBAD/mycHisA | pBR322ori; PBAD; Amp<sup>R</sup>; *araC*; C-terminal myc epitope tag. Arabinose inducible expression plasmid | Invitrogen |
| pBAD/mycHisB | pBR322ori; PBAD; Amp<sup>R</sup>; *araC*; C-terminal myc epitope tag. Arabinose inducible expression plasmid | Invitrogen |
| pKD13 | oriRSKγ Tn5 neomycin phosphotransferase (Km<sup>R</sup>), FRT, β-lactamase (Amp<sup>R</sup>) | (28) |
| pKD46 | lambda red recombinase helper plasmid | (28) |
| pSMD002 | *estP* cloned into *XhoI* site of pBAD/myc-HisA in frame with myc-His tags | this study |
| pACYC184 | p15Aori, Cm<sup>R</sup>, Tc<sup>R</sup> | (54) |
| pSMD001 | *estH* cloned into *EcoRV* site of pACYC184 | this study |
| pGPS4 | oriRSKγ Cm<sup>R</sup>, Tc<sup>R</sup> | NEB |
| pETDuet-1 | pBR322ori; *lacI*; Amp<sup>R</sup>; IPTG inducible expression plasmid | Novagen |
| pQL230 | *estP* cloned into *BglII* & *XhoI* sites of pETDuet-1 |  |
| pQL231 | *estH* cloned into *NcoI* & *EcoRI* sites of pETDuet-1 |  |
| pQL238 | *estP* cloned into *BglII* & *XhoI* sites and *estH* cloned into *NcoI* & *EcoRI* sites of pETDuet-1 |  |
| pGEX-4T1 | *lacI*, Amp<sup>R</sup>; expression plasmid for N-terminal GST fusions | GE Healthcare Life Sciences |
| pCW002<sup>a</sup> | 5047 bp GST-STh expression plasmid, Amp<sup>R</sup> | this study |
| pCW003<sup>b</sup> | *estP* cloned into *BamHI* & *EcoRI* sites of pGEX | this study |
| pYZ011<sup>c</sup> | 5047 bp GST-mSTh expression plasmid for mSTh (A14Q), Amp<sup>R</sup> | this study |
| pYZ008 | H10407 *tolC* cloned into pBAD/mycHisB | this study |
| pFLAG-CTC | 5.3 Kb plasmid, expressing C-terminal FLAG fusion proteins, Amp<sup>R</sup> | Sigma |
| pFLAG<sup>3</sup>-STh | 5609 bp Flag-Pro-STh expression plasmid, Amp<sup>R</sup> | this study |
| pSTh-FLAG<sup>3</sup> | 5612 bp Pro-STh-Flag expression plasmid, Amp<sup>R</sup> | this study |
| Km<sup>R</sup>:kanamycin resistance; Tc<sup>R</sup>: tetracycline resistance; Cm<sup>R</sup>: chloramphenicol resistance; Amp<sup>R</sup>:ampicillin resistance; NEB: New England Biolabs;<br><sup>a</sup>addgene accession number 90225: https://www.addgene.org/90225/<br><sup>b</sup>addgene accession number 90226: https://www.addgene.org/90226/<br><sup>c</sup>addgene accession number 110570: https://www.addgene.org/110570/ |  |  |

### Construction of mutants lacking production and secretion of heat stable toxins

Lamda red recombinase mediated recombination(<u>28</u>) was used to disrupt genes encoding heat stable toxins in H10407. In brief, we used the primers jf070212.1 and jf070212.2 to amplify the kanamycin cassette from pKD13 with 36 bp overhang sequence from *estH* gene, and primers of jf070212.3 and jf070212.4 to amplify kan cassatte from pKD13 with 36 bp overhang sequence from *estP* gene. The PCR product was electroporated into competent H10407 containing helper plasmid pKD46, and selected on LB plate with 50 µg/ml kanamycin. Kanamycin-resistant colonies were screened by PCR with primers of jf071612.1 and jf071612.2 for the *estH* locus, and primers of jf071612.3 and k2 for *estP*. The deletion mutants were further confirmed by toxin multiplex PCR as previously described (<u>29</u>).

To construct complemented strains, we amplified the *estP* gene with primers jf072412.3 and jf060614.1 for cloning into XhoI site of pBAD/mycHisA, yielding pSMD002; *estH* was amplified with primers of jf072414.5 and jf072414.6 and cloned into the EcoRV site of pACYC184 to produce pSMD001. We also cloned *estH* and *estP* genes individually and together in pETDuet-1 at NcoI & EcoRI sites and BglII & XhoI sites with primer pairs of jf060614.2 and jf060614.3, jf060614.4 and jf060614.5, respectively. This resulted in plasmids pQL230 (*estP*), pQL231(*estH*), and pQL238 (*estP/estH*) (<u>table 1</u>). After confirmation by sequencing, the respective plasmids were transformed into the deletion strains, and complementation confirmed by PCR. To generate an ETEC *tolC* deletion mutant, primers jf110716.44 /45 were first used to amplify a ∆tolC732:: Km^R^ fragment from JQ55031(<u>30</u>). Next, the regions flanking *tolC* in the H10407 genome were amplified with primer pairs jf110716.38/ jf110716.43 and jf110716.46/ jf110716.39 to amplify 1027 bp of upstream and 985 bp of downstream sequence, respectively. The three resulting fragments were then fused by PCR with primer pairs jf110716.38/ jf110716.39, using high-fidelity polymerase Phusion (Thermo Fisher Scientific), with denaturing for 2 min at 98°C, followed by 30 cycles of 10 s at 98°C, 30 s at 65°C, and 1.5 min at 72°C, and a final extension for 10 min at 72°C. The resulting 3,325-bp amplicon was then introduced into H10407(pKD46) as described above. Kanamycin resistant, Ampicillin sensitive colonies were then screened by colony PCR using primer pair jf112016.50/51 flanking the entire amplicon for a 4269-bp product, and primers jf101716.21/22 specific to *tolC* gene (603-bp product). Primers k2/jf112016.51 (2,024-bp product) were then used to confirm the *tolC* gene deletion and the Km^R^ cassette integration in the H10407 genome. To complement the *tolC* mutant, the *tolC* gene was amplified from H10407 genomic DNA using primers jf120716.59/jf120716.60, and the plasmid vector backbone was amplified from plasmid pBAD/myc-His B using primers jf120716.52/jf120716.53. The recombinant pYZ008 complementation plasmid was assembled using an adaptation of circular polymerase extension cloning (CPEC) (<u>31</u>) (30 s denaturation at 98°C, followed by 20 cycles of 10 s at 98°C, 30 s at 55°C, and 1.5 min at 72°C, and a final extension for 10 min at 72°C.). Following sequence verification, pYZ008 or the pBAD/myc-His B vector control plasmids were then used to transform the Δ*tolC* mutant. Complementation was confirmed by PCR with primers jf101716.21/22.

### FLAG-STH fusions

FLAG epitope fusions were constructed to introduce the 3X FLAG sequence between the signal peptide of *estH* and the beginning of the propeptide encoding region (on plasmid pFLAG^3^-STH) or at the 3’ end of *estH* (on pSTH-FLAG^3^). The 3X FLAG fragment was 1^st^ constructed by annealing complementary synthetic oligonucleotides jf092616.9 encoding + strand bases 1-43 and jf092616.10 representing– strand bases 66-19 of a 66 base pair sequence encoding the 3x FLAG peptide (DYKDHDGDYKDHDIDYKDDDDK).

To generate plasmid pFLAG^3^-STh where the 3xFLAG encoding sequence was inserted between the STh signal peptide and the STh propeptide, primers jf101916.33/ jf092616.4 were first used to amplify the nucleotides (1-63) of *estH* encoding the signal peptide. Next primers jf092616.1/ jf092616.2 were used to amplify the 3xFLAG fragment from the above synthetic oligonucleotide 3XFLAG template flanked by nucleotide extensions representing nucleotides 44-60 and 64-99 of *estH*, while primers jf092616.3/ jf101916.34 were used to amplify the 3’ end of *estH* from nucleotide 64 to the native stop codon. The three fragments were fused in a single PCR reaction using primers jf101916.33/jf101916.34. Next, the vector backbone was amplified using primers jf101916.31/ jf101916.32 from pFLAG-CTC (Sigma), followed by final assembly of pFLAG^3^-STh by CPEC. Similarly, to make plasmid pSTh-FLAG^3^ primers jf092616.5/ jf101916.35 were used to amplify 3XFLAG with a 5’ nucleotide extension representing nucleotides 197-216 of *estH*, while primers jf101916.33/jf092616.8 were used to amplify the *estH* gene with a 5’ nucleotide extension corresponding to pFLAG-CTC. The resulting amplicons were then fused in a single PCR reaction using primers jf101916.33/ jf101916.35, and assembled with the pFLAG-CTC backbone by CPEC.

### Cloning, expression, and purification of recombinant ST proteins peptides

To construct a GST-STh or mutant GST-mSTh (A14Q) expression plasmid, synthetic oligonucleotides were synthesized (IDT, Coralville, Iowa) which encompassed the region of the *estH* gene corresponding to the mature peptide minus the native start codon, preceded by an in-frame flexible linker sequence (<u>32</u>). The forward sequence (jf042715.1 for STh and jf042517.1 for mSTh), preceded by a *BamHI* overhang sequence (GATCC) and the reverse sequence (jf042715.2 for STh and jf042517.2 for mSTh), preceded by an *EcoR*I overhang sequence (AATTC) were mixed in 1:1 molar ratio, heated to 95° C for 5 minutes, and cooled to room temperate. The annealed double stranded 91 base pair DNA fragments were then cloned directly into the pGEX-4T1 vector digested with *BamH*I and *EcoR*I, yielding plasmids pCW002 and pYZ011, respectively. A similar strategy was to construct a GST-STp expression plasmid using forward oligonucleotide jf031915.5 and the reverse sequence jf031915.6, resulting in plasmid pCW003. *E. coli* TOP10(pCW002), TOP10(pCW003), and TOP10(pYZ011) were then used to express recombinant GST-STh, GST-STp, or GST-mSTh, and the resulting fusion proteins were purified by affinity chromatography. In brief, the bacterial cultures were grown in Luria broth at 37°C to an OD_600_ between 0.6 and 0.7, and induced with 1 mM IPTG final concentration for 1-3 hours. The cell pellets were re-suspended in 30 ml cold PBS containing 5 mM DTT, 1 protease inhibitor tablet (Roche), and 0.1 mg/ml lysozyme. Following sonication, supernatants were clarified by centrifugation at 12,000 rpm for 20 minutes at 4°C, followed by passage through a 0.45 µm filter. Filtered supernatants were then loaded onto columns (Glutathione Sepharose High Performance, GE Healthcare) pre-equilibrated with phosphate buffered saline (PBS), pH 7.3 (140 mM NaCl, 2.7 mM KCl,10 mM Na2 HPO4, 1.8 mM KH 2 PO4, pH 7.3). After washing with 20 column volumes of PBS containing 1 mM dithiothreitol (DTT), GST fusion proteins were eluted in fresh buffer containing (100 mM Tris-HCl, pH 8.0 and 10 mM reduced glutathione), and dialyzed against PBS.

To liberate native STh or mutant STh (mSTh) from its GST fusion partner, GST-fusion protein was dialyzed in 50 mM NH_4_HCO_3_ (pH 8.5), containing 1 unit of thrombin (Sigma) per mg GST-fusion protein at 4°C overnight, centrifuged at 4000 x g for 20 minutes a 10 kDa MWCO centrifugal filter to collect the filtrate, and dried by vacuum centrifugation. Peptide concentrations were then determined by measurement of the molar extinction coefficient at 280 nm (<u>Take3, BioTek</u>).

Plasmids pCW002, and pCW003 have been deposited in Addgene (https://www.addgene.org/) under accession numbers 90225, and 90226, respectively.

### transcriptional analysis of *estH, estP* and *tolC* genes

Confluent T84 cells were inflected with early-log phase H10407 bacteria (~10^9^cfu) for 30 or 90 min. Total RNA was isolated from adherent and planktonic (nonadherent) bacterial fractions using an RNAspin Mini Isolation Kit (GE Life Sciences, <u>25050070</u>) and treated with DNase I (ThermoFisher, <u>18068015</u>). PCR for the *arcA* housekeeping gene was used to confirm the removal of DNA. Total RNA was reverse-transcribed (SuperScript VILO cDNA Synthesis Kit, ThermoFisher, <u>11754250</u>). RNA transcripts were quantified by real-time PCR (Fast SYBR Green Master Mix, ThermoFisher, <u>4385612</u>; ViiA 7 Real-Time PCR system, <u>Applied Biosystems</u>). Primers specific to *arcA*, *estH*, *estP*, or *tolC* gene are listed in Table 2. All transcripts were normalized to *arcA*, and presented as a ratio of transcripts in adherent bacteria relative to that of planktonic bacteria.

**Table 2.**
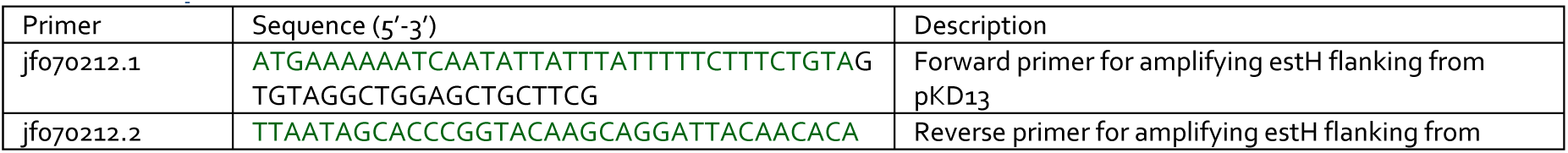
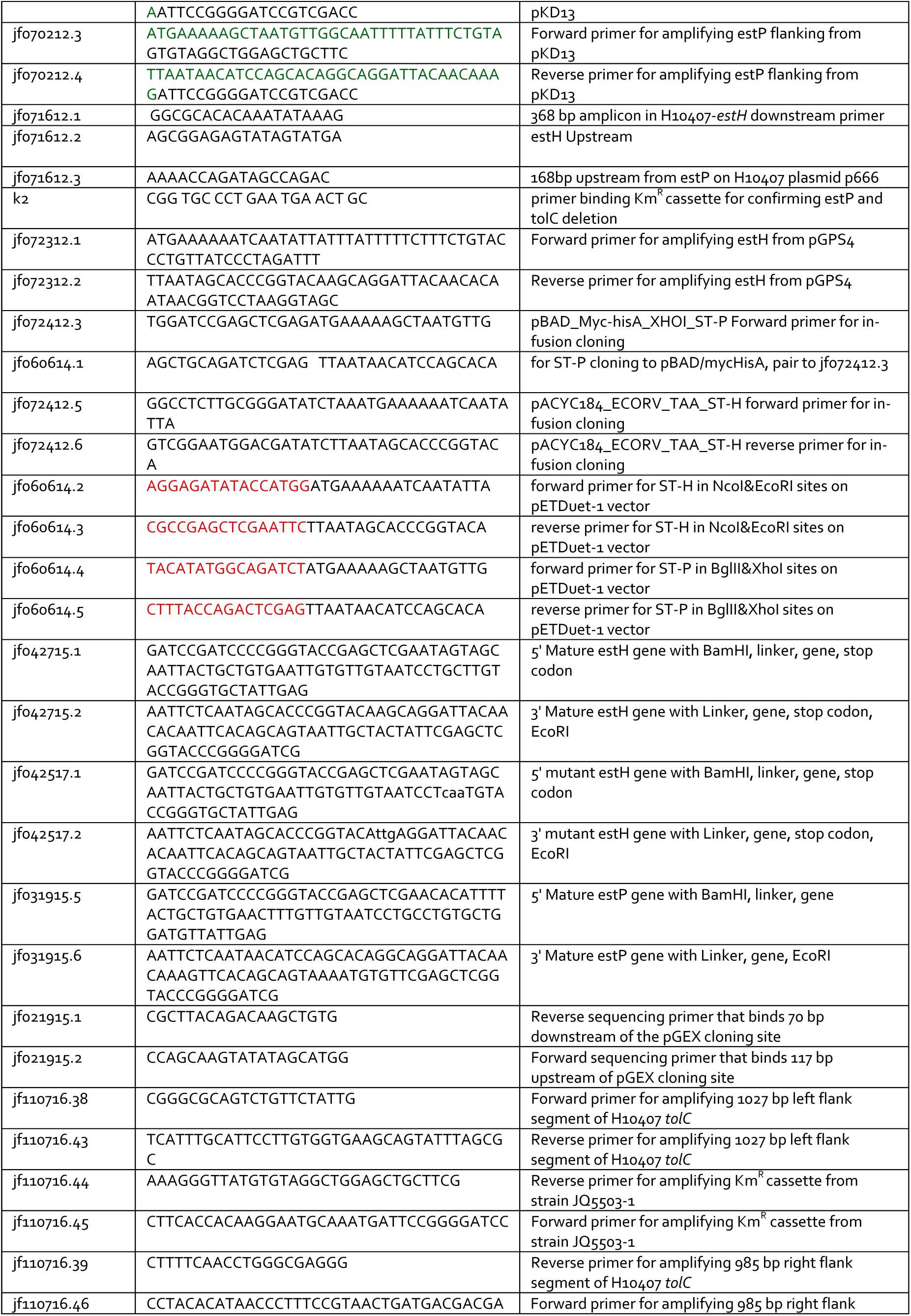
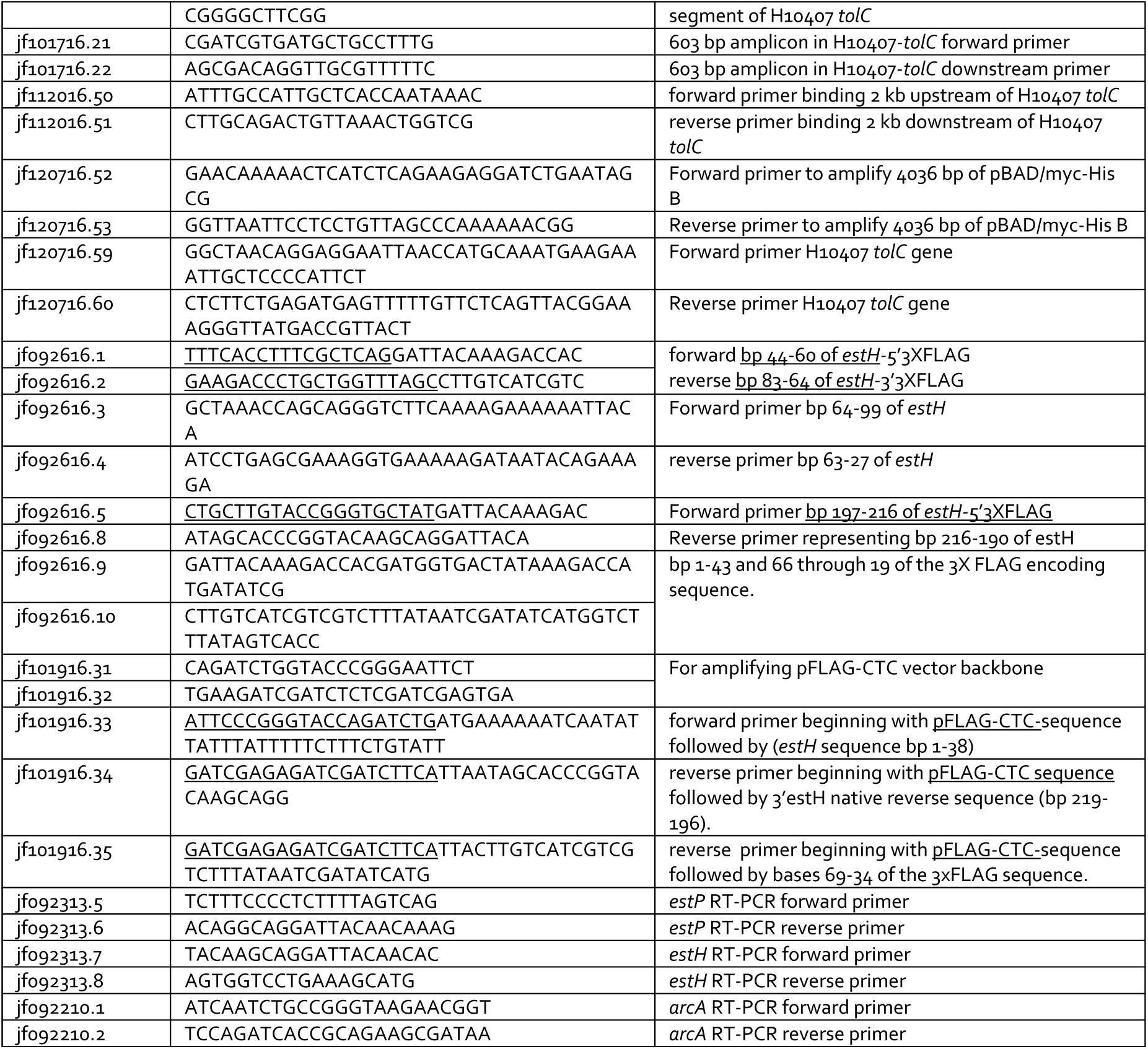
primers used in these studies

### Production and purification of anti-ST polyclonal antibody

To generate rabbit polyclonal antibody which recognizes ST-H, New Zealand white rabbits were immunized (<u>Rockland</u>) with recombinant GST-ST-H. We used lyophilized *E. coli* AAEC191A (<u>33</u>), and an immobilized *E. coli* lysate column (Pierce 44938) to absorb *E. coli*-reactive antibodies, followed by protein G column purification (HiTrap, GE <u>17-0404-01</u>). IgG GST-ST-H antibodies were affinity purified using GST-STh immobilized on nitrocellulose as previously described and anti-GST-reactive antibodies were removed by cross-absorption against GST coupled to Glutathione Agarose Resin (Gold Bio <u>G-250-100</u>, St. Louis). Anti-STp antibody, was purified from rabbit antisera (provided by Weiping Zhang, Kansas State University) by cross-absorption against an immobilized *E. coli* lysate column and then followed by affinity purification against recombinant GST-STp) immobilized on nitrocellulose membranes for affinity purification.

### Immunoprecipitation and detection of heat-stable toxins in culture supernatants

Purified anti-STh polyclonal IgG was immobilized with AminoLink Plus Coupling Resin (Pierce Direct IP kit Thermo Scientific, <u>26148</u>). Clarified culture supernatants of overnight cultures of ETEC were mixed with protease inhibitor cocktail (ThermoFisher <u>88666</u>), and filtered through a 0.45 µm filter. Sixty ml of supernatant was then filtered through 10 kDa molecular weight cutoff membrane (Amicon). Filtrates were then dried by vacuum centrifugation, and dialyzed in PBS against a 1 kDa-cutoff membrane (<u>Float-A-Lyzer G2, MWCO 0.5-1 kDa, Spectrum Labs</u>). Immunoprecipitations were conducted by incubation of filtrates with anti-STh immobilized resin for 2 hours at room temperature followed by elution with 4.5% acetic acid. Eluates were dried by under vacuum centrifugation, and reconstituted in PBS. Immunoprecipitated samples or purified protein/peptide as controls were applied directly to 0.22 µm pore-size PVDF membranes (Bio-Rad), and detected by anti-STh and/or anti-STp primary antibody and anti-Rabbit-HRP conjugated secondary antibody using Clarity Western ECL Substrate (Bio-Rad). FLAG-STh fusion peptides were prepared from strain jf2847 carrying pFLAG^3^-STh, pSTh-FLAG^3^, or the blank vector pFLAG-CTC. Briefly, following overnight growth at 37°C in lysogeny broth containing ampicillin (100 µg/ml), cultures were diluted 1:100 in fresh media, grown to OD_600_ of ~0.2, then induced with 1 mM IPTG for 7 h. 40 ml of clarified supernatant was mixed with 200 µl of anti-FLAG (M2) affinity gel (#A2220, Sigma), then incubated overnight at 4°C with agitation. After washing with TBS buffer, bound proteins were eluted with 0.1M glycine-HCl, pH 3.5, separated by SDS-PAGE, transferred to nitrocellulose for subsequent immunoblotting, and detected by anti-FLAG antibody (#F1804, Sigma).

### Confocal immunofluorescence imaging and quantification

To examine the delivery of FLAG-tagged STh toxin to intestinal epithelial cells, *estP/estH* mutant strains with or without plasmids encoding FLAG-tagged STh were grown overnight and diluted 1:50 in CAYE-ST medium, grown to OD_600_ of ~0.2, then induced with 1 mM IPTG for 2 h. The bacteria were added to T84 cells at a multiplicity of infection (MOI) of ~1:50, maintaining the induction with 1 mM IPTG. After infection with the bacteria, T84 cells were washed with PBS, fixed with 4% paraformaldehyde for 10 min at room temperature, permeabilized with 0.1% TRITON X-100 for 5 min, blocked with 1.5% BSA/PBS at 37°C for 1 h. Anti-O78 rabbit polyclonal antisera (Penn State University) diluted 1:300 in PBS with 0.02% Tween-20 (PBST) and 1.5% BSA was used to identify H10407 and monoclonal Anti-FLAG M2 antibody (#F1804, Sigma) diluted 1:500 in PBST with 1.5% BSA to detect FLAG-tagged STh. Following incubation overnight at 4°C, slides were washed 3x with PBS, and incubated for 1 hour at 37°C with goat-anti rabbit IgG (H&L) AlexaFluor-488 conjugate (ThermoFisher <u>A11070</u>) and goat-anti mouse IgG (H&L) conjugated to AlexaFluor 594 (ThermoFisher <u>A11032</u>) at a dilution of 1:500 in PBST containing 1.5% BSA. After washing, DAPI (4’, 6-diamidino-2-phenylindole) was added at 1: 6,000. Confocal microscopy images were acquired using a <u>Nikon C2+ Confocal Microscope System</u>. To quantitate binding of FLAG-epitope tagged STh molecules, fluorescence detection was normalized to the DAPI signal using NIS-Elements DUO software (v4.4).

### In vitro assessment of toxin delivery

T84 (ATCC^®^ CCL-248™) intestinal epithelial cells were maintained in DMEM/F12 (1:1) medium containing FBS (5% [vol/vol]). T84 cell monolayers were grown in 96 well plates for 24-48 hours at 37°C, 5% CO2 incubator, to > 90% confluency. Cultures of bacteria were grown overnight in lysogeny broth from frozen glycerol stocks in CAYE-ST medium. Phosphodiesterase inhibitors vardenafil hydrochloride trihydrate (#<u>Y0001647</u>), rolipram (#<u>R6520</u>), and cilostazol (#<u>C0737</u>) (all from Sigma-Aldrich), were each added to target T84 cells at a final concentration of 16.7 or 50 µM and incubated with cells for one hour. Bacteria or toxin were added to T84 monolayers seeded into 96-well plates and continued the treatment for the indicated duration. After washing in PBS cyclic GMP (cGMP) levels were determined by enzyme immunoassay (EIA) (<u>Arbor Assays, K020-H1</u>) using the acetylated protocol as directed by the manufacturer. To examine the capacity of antibodies to neutralize ST delivery, antibody against ST-H and/or EtpA was added directly to T84 cell monolayers at the indicated dilution at the time of infection. After 1.5 hours, cells were washed by PBS, and lysed, and cGMP assays were performed as described above.

## Results

### contribution of ST-H, ST-P and EAST1 to activation of cGMP in target epithelial cells

Understanding the individual contributions of ST and ST-like molecules of ETEC is relevant to development and testing of toxin neutralization strategies. H10407 encodes three peptides with the potential to activate cGMP in target intestinal epithelial cells: ST-H1, ST-P, and EAST1, a heat stable toxin originally identified in enteroaggregative *E. coli* (<u>34</u>). ST-H (ST-1b) is encoded by the *estH* gene on the large 94,797 bp p948 plasmid (<u>NCBI Genbank acccession NC_017724.1</u>). The *astA* gene which encodes EAST1 peptide (<u>35</u>) is imbedded within a IS1414 insertion sequence(<u>36</u>) is also located on the p948 plasmid immediately downstream from the *etpBAC* adhesin locus(<u>27</u>, <u>37</u>). ST-P (ST-1a) is encoded by the *estP* gene on a second 66 kb virulence plasmid <u>NCBI GenBank FN649417.1</u> in H10407.

We found that deletion of *estH* resulted in appreciable decreases in cGMP production at or near background levels of cGMP production by uninfected cells, and that complementation of *estH* in trans restored activation of this cyclic nucleotide (<u>Figure 1A</u>). In contrast, the *estP* mutant was not appreciably different than the wild type strain, and the introduction of the *estP* mutation to the *estH* strain did not yield further measurable decreases in cGMP production by target epithelial cells. However, we found that the target T84 cells used in these assays did respond to GST fusions to either ST-H or ST-P (<u>Figure 1B</u>) suggesting that these cells have the capacity to respond to either peptide. Interestingly, we found that in contrast to *estH, estP* transcription was significantly influenced by bacterial cell contact with transcription of the gene encoding ST-P enhanced by bacterial adhesion, as was the *tolC* gene which encodes the putative export channel for ST (<u>Figure 1C</u>). Deletion of *astA* gene encoding EAST on the p948 plasmid did not impact bacterial activation of cellular cGMP under the conditions of the assay (<u>Figure 1D</u>).

**Figure 1.**
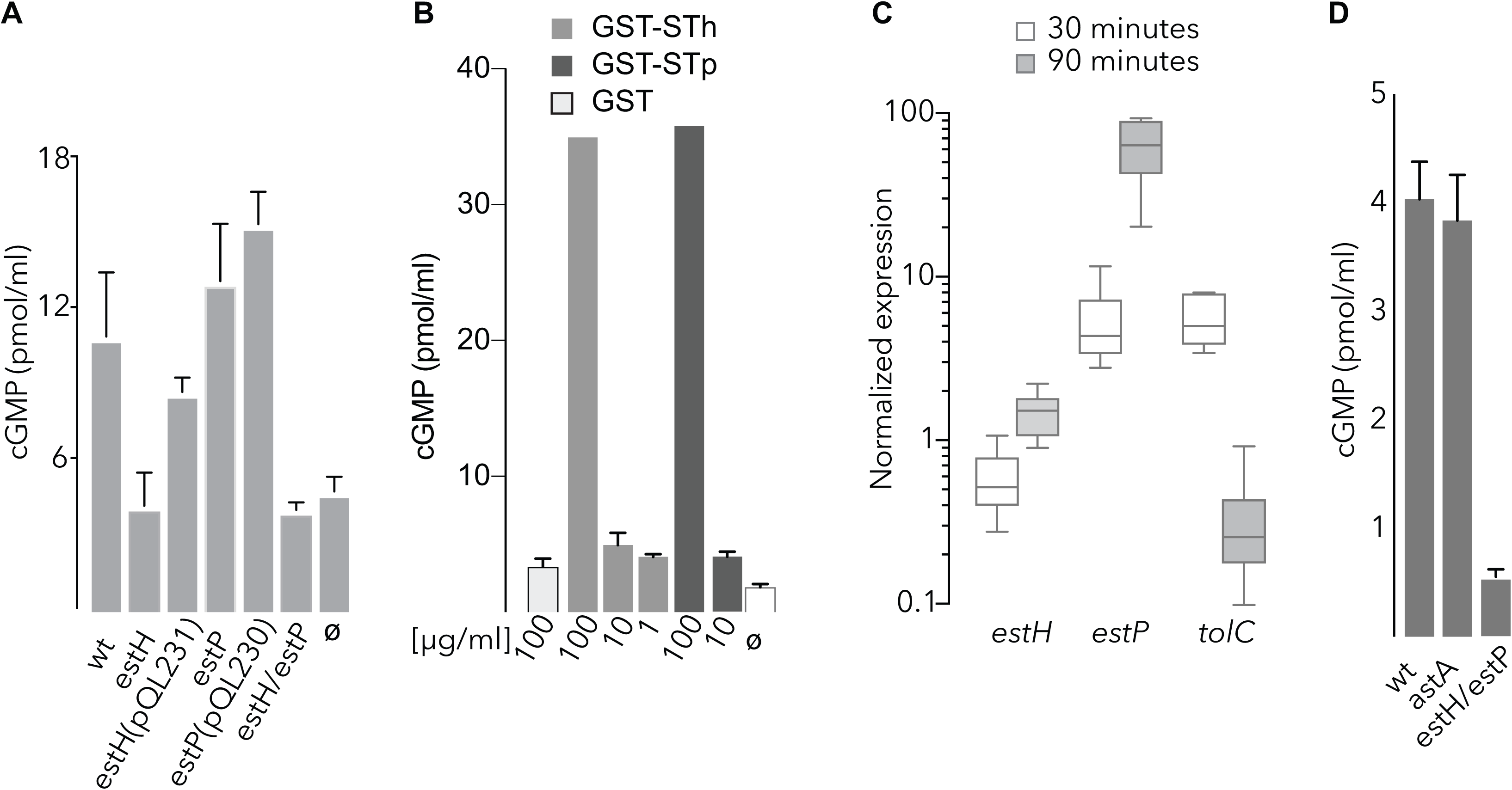
relative contribution of STh, STp, and EAST to cGMP production in target epithelia. **A.** cGMP activation of T84 target epithelial cells following infection with wild type *estH,* complemented *estH* mutant *estH*(pQL231), *estP,* the complemented *estP* mutant *estP*(pQL230), and the *estH/estP* dual deletion mutant. ø=uninfected cells. **B.** cGMP production following the addition of GST, GST-STh, or GST-STp fusions. Numbers on the x-axis represent final concentration of protein in µg/ml. ø=untreated cells. **C.** transcription of genes encoding ST toxins ST-H (*estH*), ST-P (*estP*) and TolC. Data are normalized relative to the *arcA* housekeeping gene, and represent the ratio of transcripts in attached to planktonic bacteria at 30 and 90 minutes after infection of monolayers. Whisker plots represent the range of data obtained over six replicates from two independent experiments. Horizontal lines represent mean values. **D.** Activation of cGMP in T-84 cells after addition of wild type H10407, the *astA* mutant or the dual *estH*/*estP* mutant. Data for each group represent mean of n=3 replicates ± SEM.

### TolC is required for effective ETEC secretion of ST1 toxins

The precise mechanism for secretion of heat stable toxins from ETEC strains associated with disease in humans is presently uncertain. Prior studies of heat-stable toxin investigated the secretion STb(ST-II)(<u>38</u>), or STp(STIa)(<u>39</u>) from laboratory strains of *E. coli* containing recombinant expression plasmids. While both studies suggested that the outer membrane protein TolC is involved in secreting these toxins from the recombinant *E. coli* background, there is conflicting data regarding the involvement of the STb (STII) toxin in human disease(<u>40</u>, <u>41</u>), and unlike the ST1 toxins, STb does not bind to guanylate cyclase C. Similarly, to our knowledge, there has been no verification of the role of *tolC* in mediating the secretion of either of the ST1 toxins (STh and STp) from strains of ETEC isolated from humans. Therefore, to verify the importance of the TolC in secretion of both STh (ST1b) and STp(ST1a) from ETEC which cause human illness, we constructed an isogenic *tolC* mutant in the ETEC H10407 strain, and tested the ability of the mutant bacteria to deliver ST to target epithelial cells.

We found that mutants lacking *tolC* were markedly deficient in their ability to deliver ST toxins to epithelial monolayers as we observed only background levels of cGMP production following infection with the *tolC* mutant strain, and complementation with *tolC in trans* restored the ability of the bacteria to provoke a cGMP response in targeted cells (<u>Figure 2</u>).

**Figure 2.**
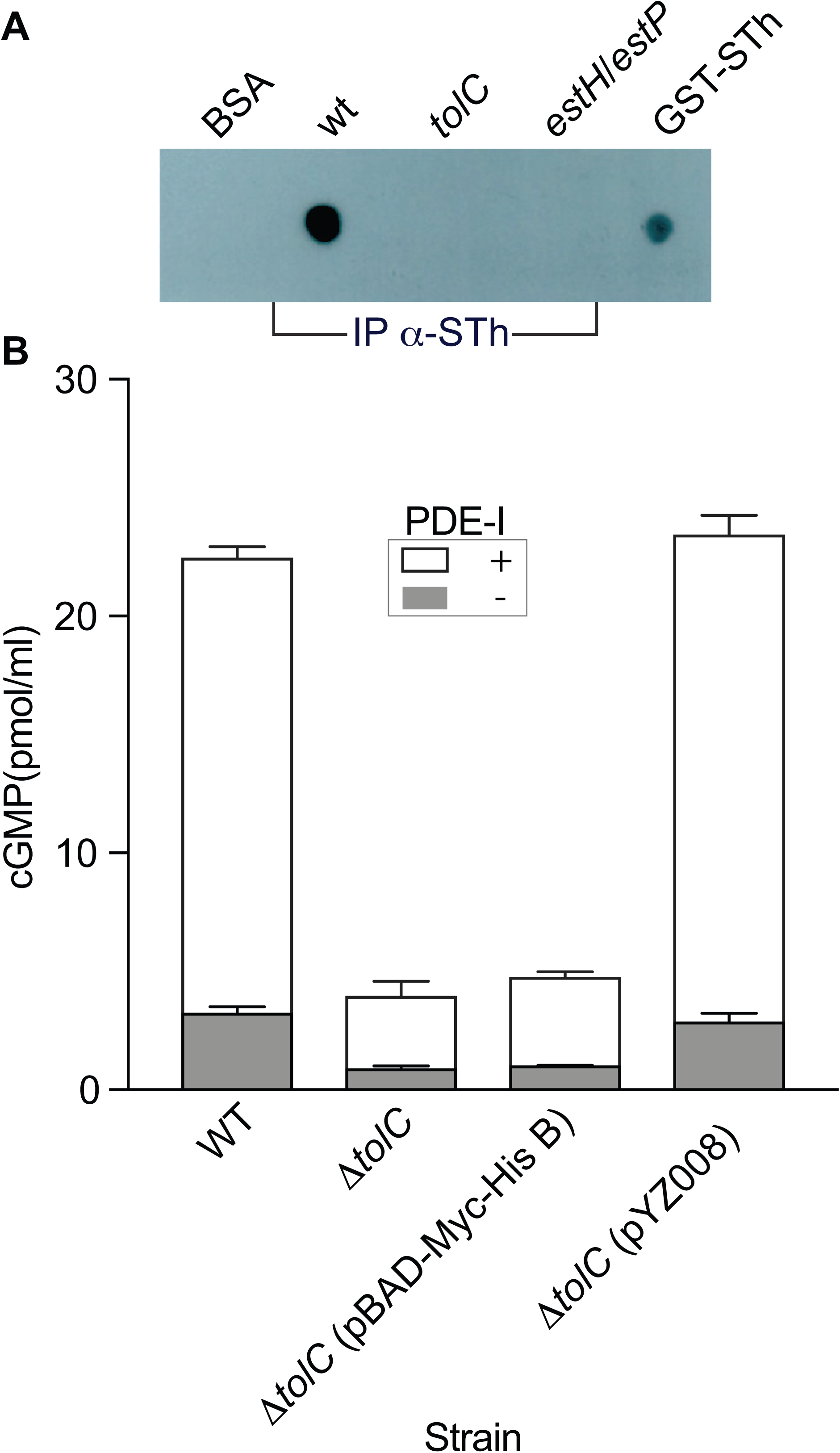
role of *tolC* in ST1 toxin secretion and delivery to target epithelial cells. **A.** Immunoblot detection of ST1 toxins in culture supernatants of the wild type H10407 strain, *tolC* mutant, or *estH*/*estP* dual deletion mutant following immunoprecipitation of culture supernatants with affinity-purified anti-STh antibodies (IP α-STh). BSA and GST-STh fusion are shown as negative and positive controls, respectively. **B.** cGMP production by T84 target epithelial cells following infection with wild type (wt) ETEC strain H10407 (ST1a, ST1b), the *tolC* mutant, the vector complemented mutant (pBAD-Myc-HisB) or the *tolC*-complemented mutant (pZY008).

### Optimal delivery of ST requires the EtpA adhesin

We have previously demonstrated that intimate interaction of ETEC with intestinal epithelial cells is essential for efficient delivery of heat labile toxin to intestinal epithelial cells(<u>42</u>). Moreover, delivery of LT requires the concerted action of several ETEC adhesins with different receptor specificities. To examine the dependence of ST delivery on bacterial adhesion we compared cGMP activation of target intestinal epithelial cells by wild type ETEC to a mutant strain lacking EtpA, a plasmid-encoded adhesin, expressed by a diverse population of ETEC(<u>43</u>). These studies demonstrated that cGMP activation in target epithelia by wild type bacteria was significantly accelerated relative to the *etpA* mutant (<u>Figure 3</u>) suggesting that efficient delivery of these small peptides also requires effective bacterial-host interactions.

**Figure 3.**
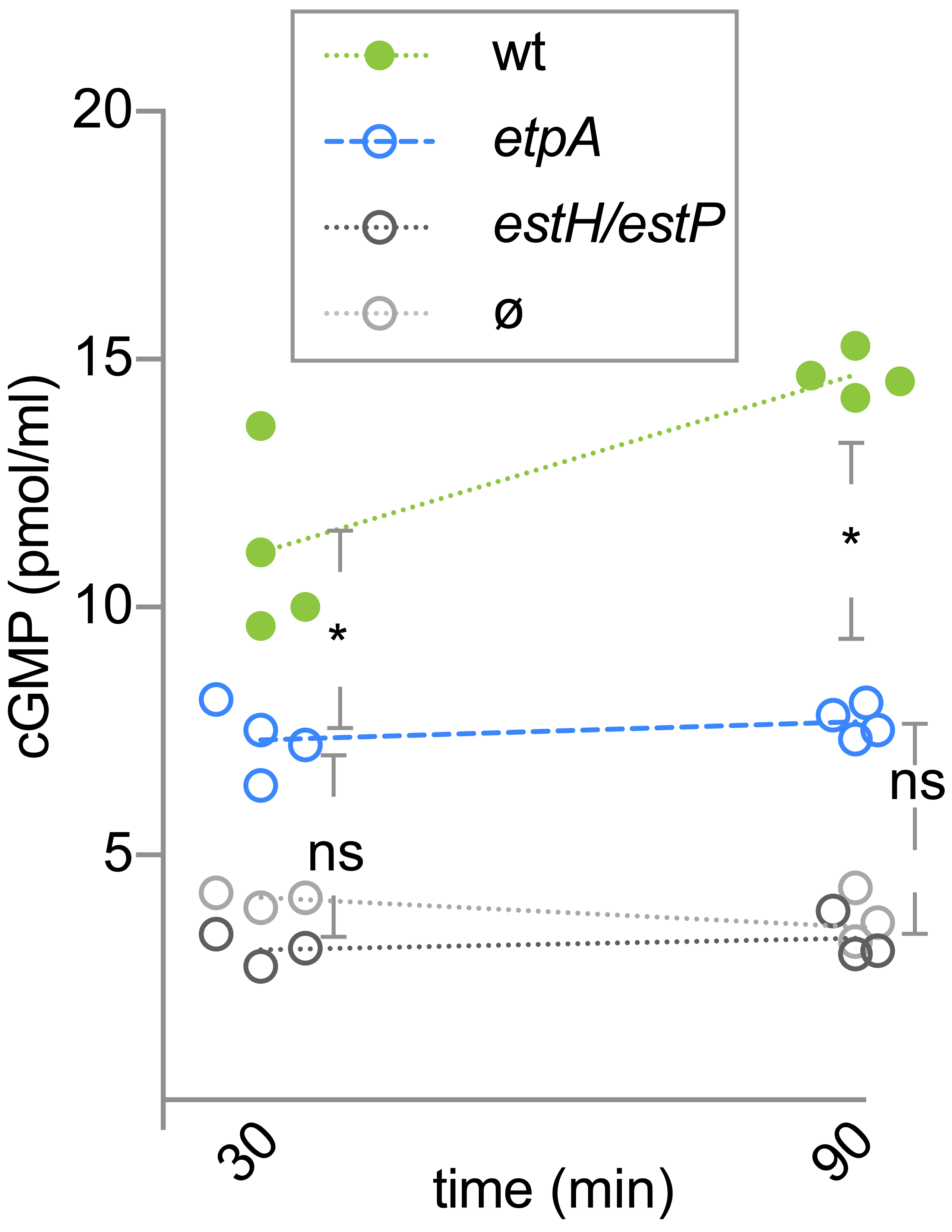
The EtpA adhesin is required for optimal delivery of heat stable toxins. Shown are comparisons of cGMP activation in target T84 intestinal epithelial cells following infection with wild type ETEC H10407, or the *etpA* mutant (n=4 replicates). The *estH/estP* dual deletion mutant and uninfected monolayers (n=3). Dashed lines in each group connect geometric mean values obtained at 30, and 60 minutes following the addition of bacteria. * represents p<0.05 obtained by Mann-Whitney, (two tailed) comparisons.

### Anti-toxin and anti-adhesin antibodies mitigate delivery of heat-stable toxins

ETEC delivery of heat labile toxin can be effectively blocked by antibodies directed against either LT or the EtpA adhesin molecule(<u>44</u>). These data suggest that anti-adhesin and anti-toxin strategies could provide complementary approaches to vaccine development. The *etpA* gene, like those encoding LT, and ST1 molecules appears to be highly conserved within the ETEC pathovar(<u>43</u>). Therefore, we examined the ability of antibodies against the EtpA adhesin to inhibit activation of cGMP in target intestinal cells.

Antibodies directed against either the EtpA adhesin molecule or STh significantly inhibited the delivery of heat-stable toxins to target cells (<u>Figure 4</u>). Although we were not able to demonstrate that the combination of these antibodies was synergistic, these data add additional support to the concept that EtpA could be useful as a target to engender coverage against a wide variety of ETEC isolates that produce heat-stable and/or heat-labile enterotoxins(<u>43</u>, <u>45</u>).

**Figure 4.**
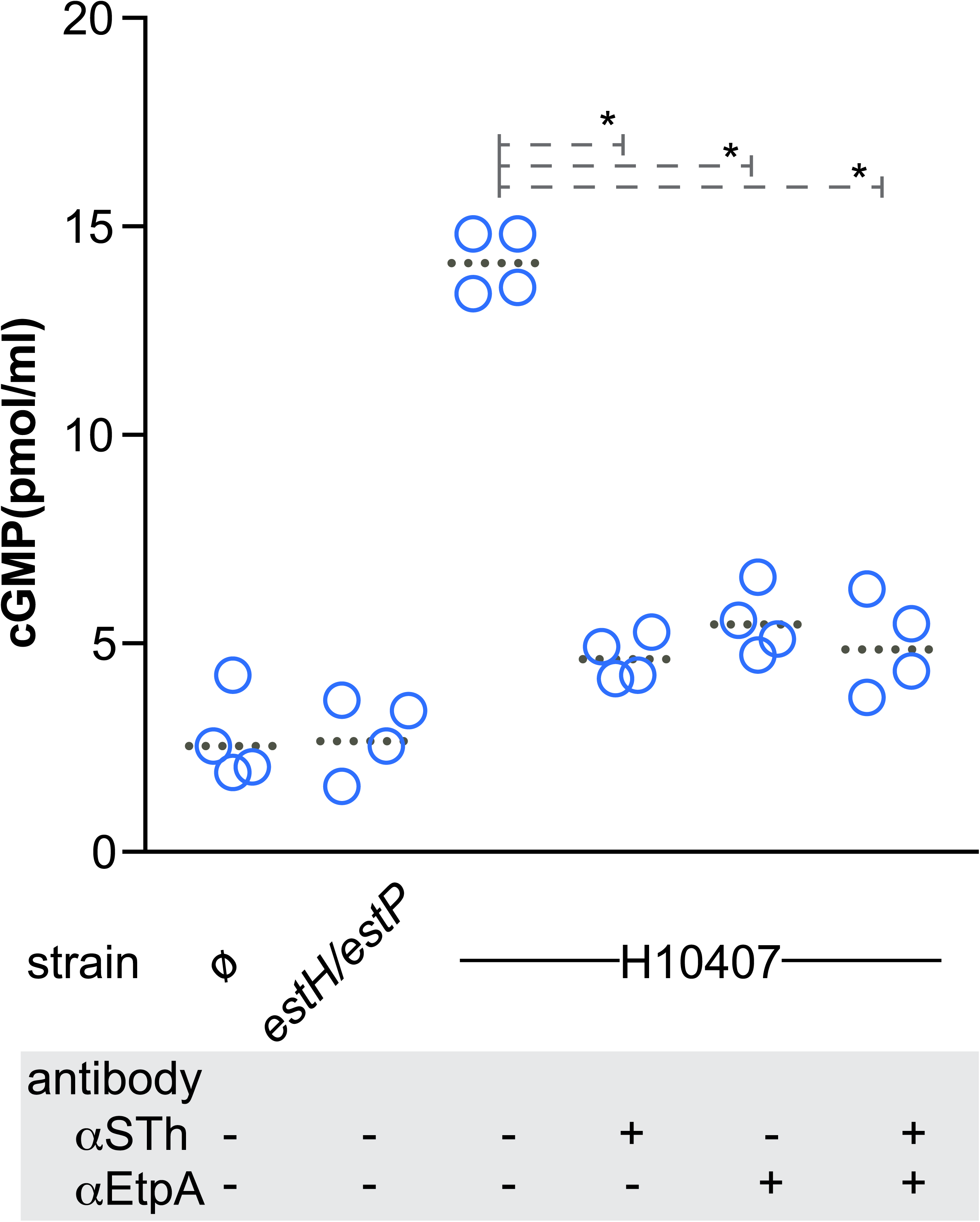
antitoxin and anti-adhesin antibodies inhibit heat-stable toxin delivery. Shown are monolayers infected with the dual heat-stable toxin mutant (*estH/estP*), wild type H10407, or uninfected control monolayers (ø). Dotted horizontal lines for each group represent geometric means. (*=0.028) by Mann Whitney two-tailed nonparametric comparisons.

### Delivery and secretion of epitope-tagged ST-H

Understanding the nature of ST secretion and its delivery to epithelial receptors could be relevant to informed development of ST toxoid molecules. ST-H is synthesized as a 72 amino acid molecule that includes a 19 amino acid signal peptide, followed by a 34 amino acid pro-peptide, and a mature ST molecule of 19 amino acids. Although most attempts to develop ST toxoids have targeted the mature peptides, there are earlier, but conflicting data regarding the precise form of ST that is secreted into the extracellular milieu (<u>15</u>, <u>46</u>), with some data suggesting that the pro-peptide may be exported with subsequent processing to the mature peptide outside the bacteria (<u>47</u>). To investigate the potential secretion of the pro-peptide, we engineered 3x-FLAG-epitope fusions to the amino terminal end of the pro-peptide region (FLAG^3^-pro-ST-H) of ST-H and compared the export, and delivery of the resulting peptides to carboxy-terminal fusions to the mature peptide (pro-ST-H-FLAG^3^, <u>schematic, figure 5</u>). We were able to recover either the amino-terminal or carboxy-terminal FLAG-tagged molecules from culture supernatants of *estH/estP* mutant ETEC transformed with the pST-FLAG^3^ or pFLAG^3^-STH plasmids, respectively (<u>figure 5a</u>). Both molecules appeared to yield functional ST mature peptides as either plasmid was sufficient to complement the ability of *estH/estP* to activate cGMP in target epithelial cells (<u>figure 5b</u>). Likewise, we were able to demonstrate binding of FLAG-epitope tagged molecules to target epithelial cells following infection with either of the complemented strains (<u>figure 5c, d</u>). Collectively, these data appear to reaffirm earlier observations of STA_3_ (ST-H)(<u>47</u>) suggesting that both the mature form (19 amino acids) of the peptide and the extended Pro-ST-H (53 amino acids) may be found outside the bacteria.

**Figure 5.**
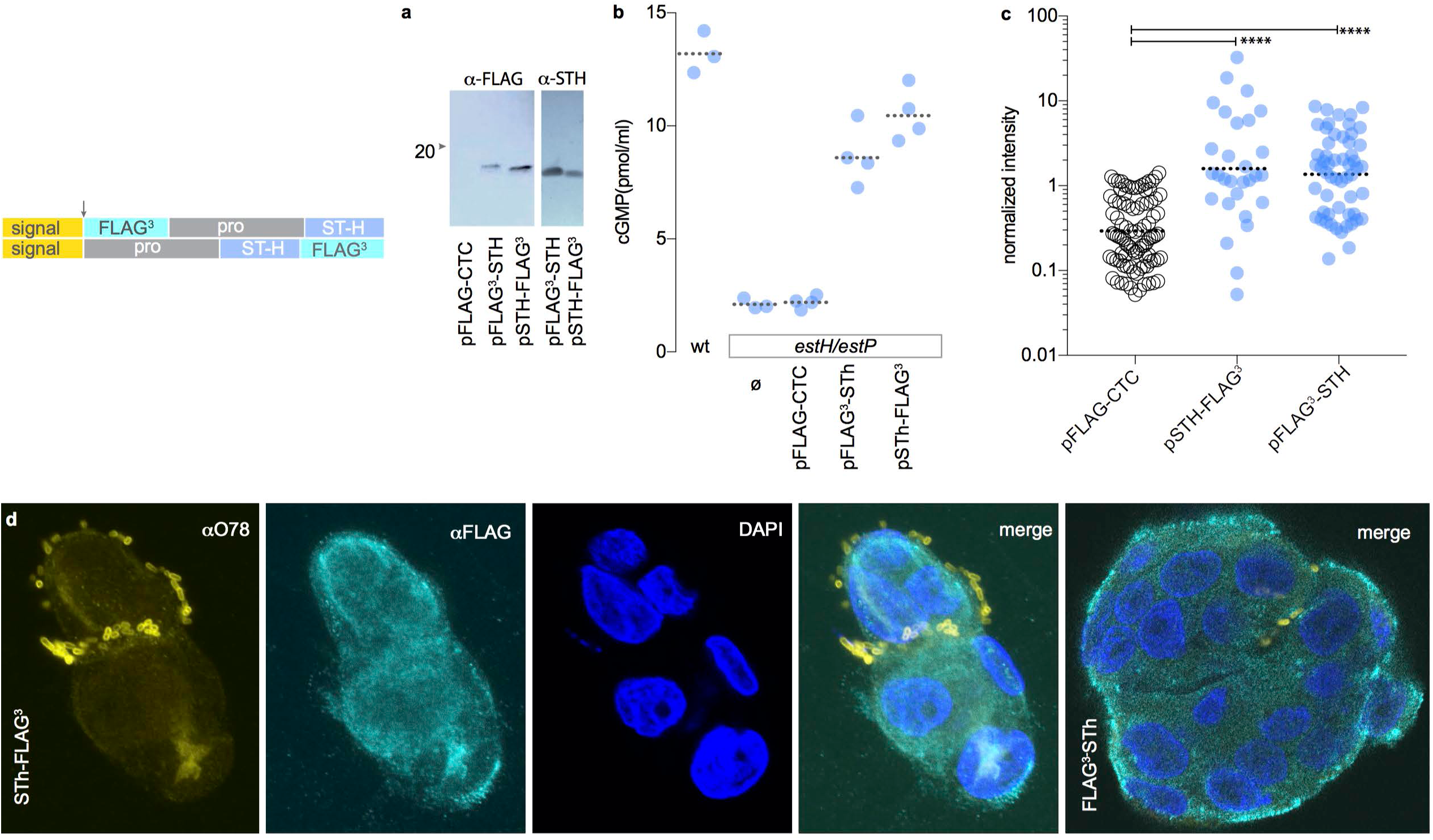
secretion and delivery of FLAG-epitope tagged ST-H. Schematic at top depicts the location of the region encoding the 3xFLAG peptide relative to the *estH* gene. The arrow at top shows the location of the predicted signal peptide cleavage site which is retained in both constructs. **a.** anti-FLAG and anti-STH immunoblots of FLAG^3^-STH and STH-FLAG^3^ peptides recovered by anti-FLAG immunoprecipitation from culture supernatants of the ETEC *estH/estP* mutant bearing the pFLAG^3^-STH and pSTH-FLAG^3^. **b.** complementation of *estP/estH* mutant strain with pSTH-FLAG^3^ or pFLAG^3^-STH restores toxicity upon infection of T84 cells. **c.** quantitation of FLAG^3^-tagged ST delivered to epithelial cells by *estH/estP* bacteria complemented with pSTH-FLAG^3^or pFLAG^3^-STH. Values represent fluorescence intensity per field. **d.** Confocal immunofluorescence images of bacteria expressing pST-FLAG^3^ (anti-O78, yellow) attached to T84 cells (nuclei, blue) and the distribution of FLAG epitope-tagged toxin (cyan).

## Discussion

Because of the significant global burden of diarrheal illness caused by ETEC, these pathogens have been a target for vaccine development since they were first identified as a cause of severe diarrheal illness more than four decades ago. Currently, there is no vaccine available that affords broad-based protection against ETEC, in part due the substantial antigenic diversity within the pathovar, and limited mechanistic insight into immunologic correlates of protection that appears to follow early infections among young children endemic areas. Moreover, many features of the molecular pathogenesis of these common pathogens have not been explored in sufficient detail to inform vaccine development.

Because ST-producing ETEC comprise a large proportion of strains associated with symptomatic diarrheal illness(<u>48</u>), we investigated the molecular contributions of the known heat-stable toxins, their proposed secretion apparatus, and bacterial adhesion to toxin delivery. The present studies suggest that efficient delivery of ST toxins is a complex process that requires the ability to directly engage host cells and at least transient adhesion afforded by EtpA and other adhesins(<u>49</u>). Moreover, as antibodies directed at either the EtpA adhesin or ST effectively impaired toxin delivery, our studies provide further support for development of a vaccine platform that combines ST-toxoid(<u>24</u>) and anti-adhesin approaches.

Interestingly, clinical studies have suggested that either ST-H originally identified in humans and ST-P can cause diarrheal illness in humans(<u>23</u>). In keeping with these observations, we found no difference in the ability of ST-H or ST-P peptides to elicit cGMP activation(<u>18</u>) in target epithelia. Although deletion of the gene encoding ST-H resulted in lower cGMP activation in infected monolayers than when the ST-P gene was deleted from the ETEC H10407 strain, we cannot exclude the possibility of a compensatory increase in ST-H secretion in the absence of a potentially competing peptide. Indeed, deletion of the *tolC* gene resulted in complete abrogation of cGMP activation in target epithelial cells, similar to deletion of both of the ST1 toxin genes from H10407.

Earlier studies also noted that many ETEC strains including H10407 bear copies of an insertion sequence(<u>36</u>) that encompass the *astA* gene encoding EAST1(<u>35</u>), a cGMP activating peptide structurally similar to ST1 that was originally identified as a heat-stable enterotoxin in enteroaggregative *E. coli(<u>34</u>).* However, we saw no appreciable decrease in cGMP in target epithelial cells following infection with the *astA* mutant, compared to wild type ETEC. We cannot rule out the possibility of EAST1 expression from additional copies of *astA* not residing on the 92 MDa virulence plasmid, or that EAST1 is not optimally expressed in *vitro*. However, the complete absence of a cGMP response in cells infected with the ST-1a/ST-1b deletion mutant might alternatively suggest that EAST1 does not contribute to ETEC virulence and that further toxoid vaccine development can simply focus on engendering neutralizing antibody responses to the established ST1 and LT enterotoxins.

The reaffirmation of TolC as a key mechanism for export of ST1 toxins could be relevant to management of diarrheal illness. The rapid emergence of multi-drug resistance in the Enterobacteriaciae that is in part dependent on drug efflux through TolC has stimulated interest in efflux inhibitors to enhance the potency of available antimicrobial agents(<u>50–52</u>). Theoretically, these inhibitors could offer novel therapeutic agents for treatment of ETEC. Our studies also revisit the concept that the larger pro-peptide form of ST-H may be exported by the bacteria, and the data presented here are consistent with prior observations suggesting that some processing of the pro-peptide occurs outside the bacteria(<u>15</u>, <u>47</u>). Further study will be needed however to determine whether the pro-peptide sequence contributes to immunologic recognition of ST following infection and whether these larger molecules might be exploited in the development of improved toxoids to neutralize ST.

## Acknowledgements

These studies were supported in part by funds from the National Institute for Allergy and Infectious Diseases (NIAID) grants R01AI89894, R01AI126887; and The Department of Veterans Affairs (5I01BX001469-05); CW was supported in part by a Washington University/HHMI Summer Undergraduate Research Fellowship (HHMI SURF).

